# Noncoding regulatory mutations contribute to the aberrant gene expression profile of neuroblastomas

**DOI:** 10.64898/2026.08.10.743941

**Authors:** Brianna Jones, Gaurav Seth, Abby Robertson, Arko Sen

## Abstract

Comprehensive analyses of whole-genome and exome sequencing data from high-risk neuroblastoma tumors have identified relatively few recurrent and clinically actionable protein-coding driver mutations at initial diagnosis. This suggests that noncoding genetic variations that alter regulatory sequences and impact gene expression are important drivers of neuroblastoma tumorigenesis. Allele-specific expression (ASE), which quantifies differences in expression between the two alleles of a gene, is a powerful approach for identifying genes with altered dosage resulting from cis-regulatory variation. In this study, we compared ASE profiles of 156 neuroblastoma patients with 220 adrenal gland tissues from the Genotype-Tissue Expression (GTEx) Project to identify 1,363 neuroblastoma-specific ASE genes. We demonstrated that these genes were enriched for biological processes relevant to neuroblastoma pathogenesis including chromatin remodeling, protein ubiquitination, and sympathetic nervous system development, and were preferentially associated with the adrenergic transcriptional program. To elucidate the genetic mechanism underlying aberrant expression of neuroblastoma-specific ASE genes we integrated our ASE data with somatic copy number alterations (SCNA) and somatic mutation profiles. We observed that although many neuroblastoma specific ASE genes were associated with chromosomal gains and losses, a substantial subset was located within copy-number-neutral genomic regions. These genes showed significant enrichment for noncoding somatic single-nucleotide variants (SNVs) within intronic and distal intergenic neuroblastoma-specific open chromatin regions. Furthermore, functional analyses revealed that many of these SNVs disrupt transcription factor binding sites for GATA3, a core component of the regulatory circuitry that maintains the adrenergic identity of neuroblastoma cells, and neuroblastoma-specific ASE genes harboring such mutations were significantly more likely to belong to the GATA3 regulon than expected by chance. Together, our findings identify novel molecular targets of noncoding regulatory mutations in neuroblastoma and highlight the important yet underappreciated contribution of noncoding genetic variation to neuroblastoma tumorigenesis.

## INTRODUCTION

Neuroblastoma is the most common type of extracranial solid tumor in children that occurs in the adrenal medulla and along the paraspinal ganglia. It represents 8% of all childhood cancers but 15% of childhood cancer mortality in the United States [1]. Comprehensive exome sequencing of high-risk tumors has identified a few protein-coding driver mutations [2]. These include activating mutations in the ALK tyrosine kinase and missense or truncating mutations in the chromatin remodeler ATRX [2–4]. However, beyond these alterations, pathogenic protein-coding mutations in primary neuroblastomas are exceedingly rare.

Instead, the neuroblastoma genome is characterized by somatic copy number alterations (SCNA), which are associated with adverse clinical outcomes. The most frequent SCNA is focal amplification of chromosome 2p24, encompassing the MYCN oncogene. This amplification occurs in 20–30% of patients and is a defining feature of high-risk disease [5, 6]. Other recurrent SCNAs include arm-level hemizygous deletions of chromosomes 1p, 3p, and 11q, as well as gain of an unbalanced gain chromosome 17q, all of which are predictive of poor patient survival [7–9].

Allele-specific expression (ASE) measures differences in expression between the two alleles of a gene. It can be applied to a single sample and is insensitive to environmental and trans-acting factors. These features make ASE a powerful tool for identifying genes with altered expression dosage due to cis-acting regulatory variations [10]. We have developed an analytical framework that quantifies gene-level ASE by combining allelic imbalance information across multiple heterozygous variants mapping to gene exons [11]. Using this algorithm, we have demonstrated that ASE in neuroblastomas is frequently associated with SCNA; however, many genes, including genes residing within regions affected by SCNA, also exhibit ASE in copy-neutral tumors. These observations suggest that additional, as-yet-unidentified genetic variations contribute to gene expression dysregulation in neuroblastomas [11].

Emerging evidence implicates noncoding single-nucleotide variants (SNVs) that disrupt regulatory elements such as promoters and enhancers as important contributors to transcriptional dysregulation in neuroblastomas. For example, a germline G to T transversion within the first intron of LMO1 disrupts a GATA binding site in its super-enhancer region, leading to reduced LMO1 expression, loss of the adrenergic cell state, and decreased tumor penetrance in vivo [12]. Beyond germline variations, noncoding somatic mutations have also been implicated in gene expression dysregulation in neuroblastomas. Integrated analysis of WGS data from patient samples and DNA Hypersensitive Sites (DHS) from neuroblastoma cell lines has identified several mutations within neuroblastoma-specific open chromatin regions. These mutations have higher pathogenicity scores than mutations within non-specific open chromatin regions, and they commonly disrupt transcription factor binding sites (TFBS) for transcription factors (TFs) involved in cell-cycle phase transitions, cell differentiation, and immune-related pathways [13–15].

However, despite recent advances, studies interrogating noncoding somatic mutations in neuroblastomas remain limited. This is likely due to their low recurrence across patient populations, which necessitates large sample sizes for detection. Expanding patient cohorts for rare pediatric cancers like neuroblastoma can be costly and logistically challenging. Furthermore, it is difficult to characterize their impact on gene expression because regulatory elements are often spread across large genomic regions and different regulatory elements may be mutated in different patients. For these reasons, aggregation-based methods designed to discover protein-coding driver mutations are ineffective for noncoding somatic mutations, highlighting the need for alternative strategies.

In this study, we present an integrative framework for identifying noncoding regulatory mutations and their target genes in neuroblastoma. First, we define neuroblastoma-specific allele-specific expression (NB-ASE) genes by comparing the frequency of ASE events between neuroblastoma tumors and normal adrenal gland tissues. Next, we evaluate whether noncoding somatic mutations are enriched within the regulatory regions associated with NB-ASE genes. Finally, we functionally characterize these mutations by identifying the TFBS near NB-ASE genes they disrupt and determining whether the affected NB-ASE genes are members of the corresponding TF’s regulons.

Applying this framework to our dataset, we have made several key discoveries. We have identified 1,363 NB-ASE genes and found that they are enriched for biological processes relevant to neuroblastoma pathogenesis, including chromatin remodeling, protein ubiquitination, and sympathetic nervous system development, and are preferentially associated with the adrenergic transcriptional program. We have demonstrated that noncoding somatic mutations are enriched within neuroblastoma-specific open chromatin regions located in distal intergenic and intronic regions of NB-ASE genes. Finally, we discovered that these noncoding mutations most frequently affect the regulons of TFs such as GATA3, GABPA, ZBTB33, and JUND and are associated with the expression dysregulation of well-characterized epigenetic regulators, including KMT2C (MLL3), in neuroblastoma.

## RESULTS

### Identifying genes that show neuroblastoma-specific ASE patterns and determining their role in neuroblastoma tumorigenesis

ASE quantifies differences in expression between the two alleles of a gene and is a powerful approach for identifying genes with altered dosage resulting from cis-acting regulatory variation. To determine the ASE profile of neuroblastoma patients, we analyzed matched whole-genome sequencing (WGS) and RNA sequencing (RNA-seq) data from 73 patients in the Pediatric Cancer Genome Project (PCGP) cohort and 83 patients in the Kids First Pediatric Research Program (Kids First) cohort. To estimate ASE, we identified heterozygous sites located within gene exons and retained sites supported by at least 10 RNA-seq reads. Then we applied our previously published statistical framework, which integrates allelic imbalance information across multiple heterozygous exonic variants while accounting for technical sources of variation, including genotyping errors, sequencing errors, and overdispersion of RNA-seq read counts, to estimate gene-level ASE [11].

In the PCGP cohort, we were able to test an average of 9,069 genes for ASE, and at a false discovery rate (FDR) threshold of 0.1 or 10% we identified an average of 402 significant ASE genes, representing approximately 4.4% of testable genes per sample (Supplemental Figure S1A & B). In the Kids First cohort, we were able to test an average of 5,437 genes per sample and identified an average of 113 significant ASE genes per tumor, corresponding to 2.2% of testable genes (Supplemental Figure S1A & B).

Because tumor sequencing datasets typically contain both malignant tumor cells and normal stromal cells, contamination by nonmalignant cells may impact our ability to detect ASE events. To examine this possibility, we used FACETS (Fraction and Allele-Specific Copy Number Estimates from Tumor Sequencing) to estimate allele-specific copy number states, tumor purity, and tumor ploidy for each tumor sample [16]. We then plotted the proportion of genes exhibiting significant ASE (FDR ≤ 0.1) against FACETS-derived tumor purity estimates across the cohort. No significant association was observed between tumor purity and the proportion of ASE genes detected (Pearson’s r = -0.071, P = 0.38), indicating that ASE detection was largely robust to variation in tumor purity within this dataset (Supplementary Figure 1D). This robustness likely reflects the fact that ASE is quantified as a within-sample allelic ratio normalized by local read depth, rather than an absolute expression measure which is sensitive to bulk tumor content. Based on these quality-control analyses, we decided to exclude samples with fewer than 5,000 testable genes for ASE or a tumor purity estimate below 30% from downstream analysis resulting in a final study cohort of 112 primary neuroblastoma tumors.

Recurrent ASE can arise from several nonpathogenic mechanisms, including common germline polymorphisms, genomic imprinting, and random monoallelic expression [10, 17, 18]. To distinguish allelic imbalance caused by these processes from imbalance driven by somatic alterations, we compared ASE profiles between neuroblastoma tumors and normal adrenal gland tissues from the Genotype Tissue Expression (GTEx) project. Although the GTEx adrenal gland samples were derived from adults, our previous study demonstrated that they provide an effective reference for identifying neuroblastoma-specific ASE (NB-ASE) genes [11]. Therefore, similar to our published study, we defined NB-ASE genes as genes that were testable for ASE in at least 10 neuroblastoma samples and 10 adrenal gland samples and exhibited significant ASE (FDR ≤ 0.1 or 10%) in at least 3 neuroblastoma samples but in no more than 1 adrenal gland sample resulting in the identification of 1,363 candidate NB-ASE genes (Supplemental Table S4).

To confirm that this simple hard filtering approach effectively isolates genes relevant for neuroblastoma pathogenesis, we examined the ASE profiles of imprinted genes and neuroblastoma tumor suppressor genes. We reasoned that imprinted genes should exhibit recurrent ASE in both normal adrenal gland and neuroblastoma samples, whereas tumor suppressor genes affected by neuroblastoma-specific genetic lesions should primarily display ASE in tumor tissues. Consistent with this expectation, imprinted genes such as MEST [19] exhibited frequent ASE in both adrenal gland and neuroblastoma samples, while established neuroblastoma tumor suppressors including KIF1B, CHD5, and NF1 displayed recurrent ASE predominantly in tumor samples [11, 20–22]. These findings recapitulate observations from our previous study and support the use of normal adult adrenal gland tissues as a reference for identifying NB-ASE events (Figure 1A).

**Figure 1.**
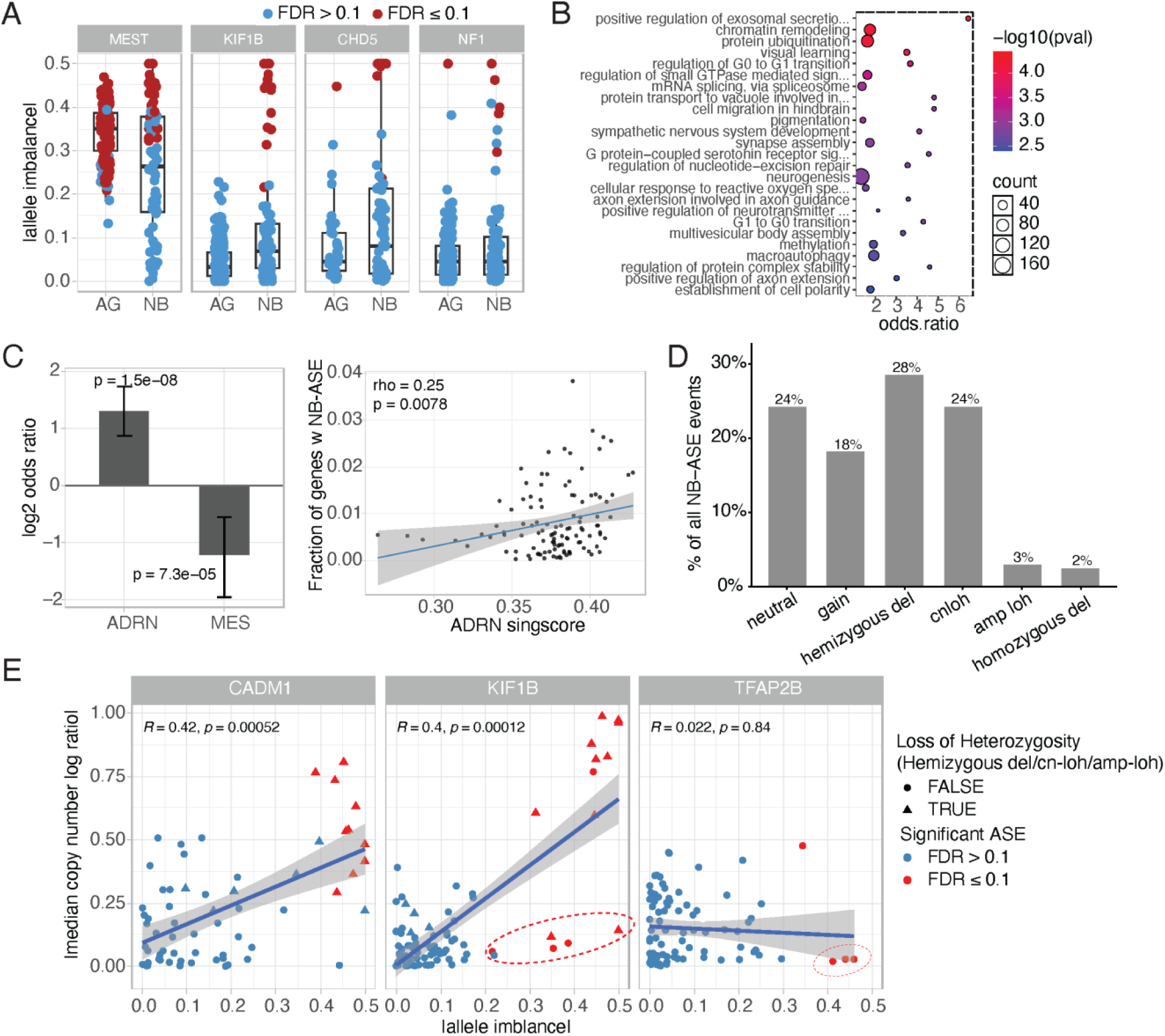
Neuroblastoma-specific allele-specific expression (NB-ASE) genes are enriched for regulatory and developmental processes linked to neuroblastoma cell identity and include a minority of genes with allelic imbalance patterns not explained by somatic copy-number alteration. A) Absolute allelic imbalance (|allele imbalance|) for the imprinted gene MEST and the neuroblastoma tumor suppressor genes KIF1B, CHD5, and NF1 in normal adrenal gland (AG) and neuroblastoma (NB) samples. Each point represents one sample; points are colored by ASE significance (FDR > 0.1, blue; FDR ≤ 0.1, red). B) Gene Ontology (Biological Process) enrichment for NB-ASE genes, using all genes showing ASE as the background universe. The top 25 enriched terms are shown, ranked by odds ratio; dot size indicates the number of NB-ASE genes annotated to each term, and dot color indicates statistical significance (−log10 P, weight01 Fisher’s exact test). C) *Left*, enrichment of NB-ASE genes among adrenergic (ADRN) and mesenchymal (MES) neuroblastoma signature genes, expressed as log2 odds ratio (Fisher’s exact test; error bars, 95% CI). *Right*, per-tumor correlation between ADRN signature score (singscore) and the fraction of testable genes exhibiting NB-ASE (Spearman’s ρ = 0.25, P = 0.0078); each point represents one tumor, with a linear fit (shaded region, 95% CI). D) Depicts the percentage of all NB-ASE events across the cohort stratified based on copy number status. E) Relationship between local copy-number ratio (lcnr, median) and |allele imbalance| for three representative genes — CADM1, KIF1B, and TFAP2B — across tumors. Points are colored by ASE significance (FDR ≤ 0.1: red; FDR > 0.1: blue) and shaped by loss of heterozygosity (LOH) status of overlapping genomic region. Pearson’s R and P-value are shown for each gene. Red circles highlight samples with significant allelic imbalance despite a copy-neutral state, illustrating SCNA-independent ASE events at KIF1B and TFAP2B.

To determine the disease relevance of NB-ASE genes, we performed Gene Ontology (GO) enrichment analysis for biological processes using all genes tested for ASE (FDR ≤ 0.1) as the background universe. GO terms were evaluated using the “*weight01*” algorithm, which accounts for the hierarchical structure of the GO graph and dependencies among related terms. A minimum node size of 10 genes was applied to exclude poorly powered categories. We then examined the top 25 enriched biological processes associated with NB-ASE genes and observed that they were significantly enriched for biological processes relevant to neuroblastoma pathogenesis, including chromatin remodeling (GO:0006338, P = 6.5 × 10⁻^5^), protein ubiquitination (GO:0016567, P = 7.3 × 10⁻^5^), and sympathetic nervous system development (GO:0048485, P = 0.00103) (Figure 1B).

Finally, to further illustrate the biological relevance of NB-ASE genes, we examined their association with neuroblastoma cell states. The adrenergic (ADRN) and mesenchymal (MES) cell states represent the two core transcriptional identities and states of neuroblastoma cells [23]. We reasoned if NB-ASE genes largely reflected technical noise — such as mapping artifacts, coverage biases, or transcriptional variability — their distribution across these two programs would be expected to be proportional with no preferential association with either cell state. In contrast, a skewed distribution of NB-ASE genes between the ADRN and MES transcriptional programs would argue against a purely technical origin.

To determine whether NB-ASE genes were non-randomly distributed with respect to established neuroblastoma cell states, we first tested for overlap between NB-ASE genes and published adrenergic (ADRN; n = 369) and mesenchymal (MES; n = 485) signature gene sets using all ASE genes in the cohort as the background universe. ADRN and MES signature genes were obtained from van Groningen et al [23]. Fisher’s exact test revealed a significant enrichment of ADRN signature genes among NB-ASE genes (4.5% of ADRN genes vs. 1.8% of background genes; OR = 2.46, 95% CI = 1.81-3.30, adjusted P = 3.01 × 10^-8^) (Figure 1C, *left panel*). In contrast, MES signature genes were significantly depleted among NB-ASE genes (1.4% vs. 3.32%; OR = 0.43, 95% CI = 0.26-0.68, P = 7.27 × 10^-5^) (Figure 1C, *left panel*). These findings indicate that genes exhibiting NB-ASE patterns are preferentially associated with the adrenergic cell state.

To further validate this relationship, we examined whether the burden of NB-ASE events (defined as number of NB-ASE events divided by the total number of ASE events) was associated with ADRN or MES transcriptional signatures across tumor samples. For each tumor, ADRN and MES signature scores were calculated using singscore (v1.30.0) (RRID:SCR_028057) [24], and associations with NB-ASE burden were assessed using Spearman’s rank correlation. We observed that the ADRN scores were significantly positively correlated with NB-ASE burden (Spearman’s ρ = 0.25, adjusted P = 0.015) (Figure. 1C, *right panel*), whereas MES scores showed no significant association (ρ = 0.11, adjusted P = 0.23). These results suggest that tumors with a more adrenergic transcriptional state show more frequent ASE of NB-ASE genes, further supporting their relevance to neuroblastoma pathogenesis.

### A subset of NB-ASE genes shows allelic imbalance independent of local copy-number alteration

Next, we examined the genetic mechanism that contributes to ASE profile of NB-ASE genes. Our previous study had demonstrated that a majority of ASE signal in neuroblastoma are attributable to SCNA [11]. To determine the extent to which recurrent allelic imbalance at NB-ASE genes is attributable to SCNA in PCGP and Kids First cohort, we intersected significant ASE calls (FDR ≤ 0.1) for NB-ASE genes with allele-specific copy-number segments derived from FACETS for every neuroblastoma tumor. This analysis yielded 7,520 classifiable NB-ASE events across 112 tumors, spanning 1,363 NB-ASE genes. We then calculated the percentage of NB-ASE events stratified by FACETS copy-number segment labels. We observed that while 76% of events were attributable to SCNA, 24% of NB-ASE events (mapping to 691 NB-ASE genes) were located within copy neutral regions. Thus, a substantial fraction of gene dysregulation in neuroblastoma is driven by SCNA independent cis-regulatory mechanisms.

To examine this relationship at the level of individual loci, we compared the absolute allele imbalance (|allele imbalance|) for genes with the absolute log2 copy-number ratio (|cnlr median|) of the overlapping genome segment across tumors for three representative genes implicated in neuroblastoma pathogenesis CADM1, KIF1B, and TFAP2B (Figure 1E) [20, 25–27]. CADM1 (n = 63 tumor-gene observations; Spearman’s ρ = 0.42, P = 0.00052) and KIF1B (n = 86; ρ = 0.40, P = 0.00012) showed a significant positive correlation between copy-number magnitude and allelic imbalance, consistent with SCNA as a major contributor to ASE at these loci. However, KIF1B also included tumors with significant allelic imbalance in copy neutral samples, consistent with additional SCNA independent mechanisms underlying gene dysregulation. In contrast, TFAP2B (n = 84) showed no relationship between copy-number magnitude and allelic imbalance (ρ = 0.022, P = 0.84), with tumors mainly exhibiting significant ASE in copy neutral samples, indicating that allelic imbalance at this locus is largely SCNA-independent. Based on these observations, we hypothesized that allele imbalance of a significant fraction of copy neutral NB-ASE genes is driven in part by noncoding somatic mutations that impact promoters and enhancers.

### Noncoding somatic mutations are enriched within putative regulatory regions of NB-ASE genes

To determine whether noncoding mutations contribute to the ASE profile of NB-ASE genes, we examined whether they are enriched near NB-ASE genes compared to non-NB-ASE genes (i.e., genes showing ASE in both neuroblastoma and adrenal gland tissue). Prior to enrichment analysis, we removed SNVs covered by fewer than 10 WGS reads (DP < 10) and SNVs located within low-mappability (repeats), exonic, or splicing-associated genomic regions. To control for the potential confounding effect of SCNA, we also restricted the analysis to SNVs and ASE genes residing within copy-neutral regions in any given tumor. We then constructed, for each sample, annotations linking each noncoding SNV to the nearest NB-ASE gene and separately to the nearest non-NB-ASE gene with a TSS within ±1 Mb, ±0.5 Mb, or ±0.25 Mb of the SNV. Then for each sample and gene class, we calculated the noncoding SNV rate per gene (number of unique noncoding SNVs divided by number of genes). Each tumor contributes one rate per class, and only tumors with both classes represented were retained (n = 86, 85, and 83 for the ±1 Mb, ±0.5 Mb, and ±0.25 Mb windows, respectively). Comparing log1p-transformed rates we found that the rate of somatic SNVs was consistently higher for NB-ASE than non-NB-ASE genes across all distance thresholds, suggesting that noncoding SNVs are enriched near NB-ASE genes (Figure 2A).

**Figure 2.**
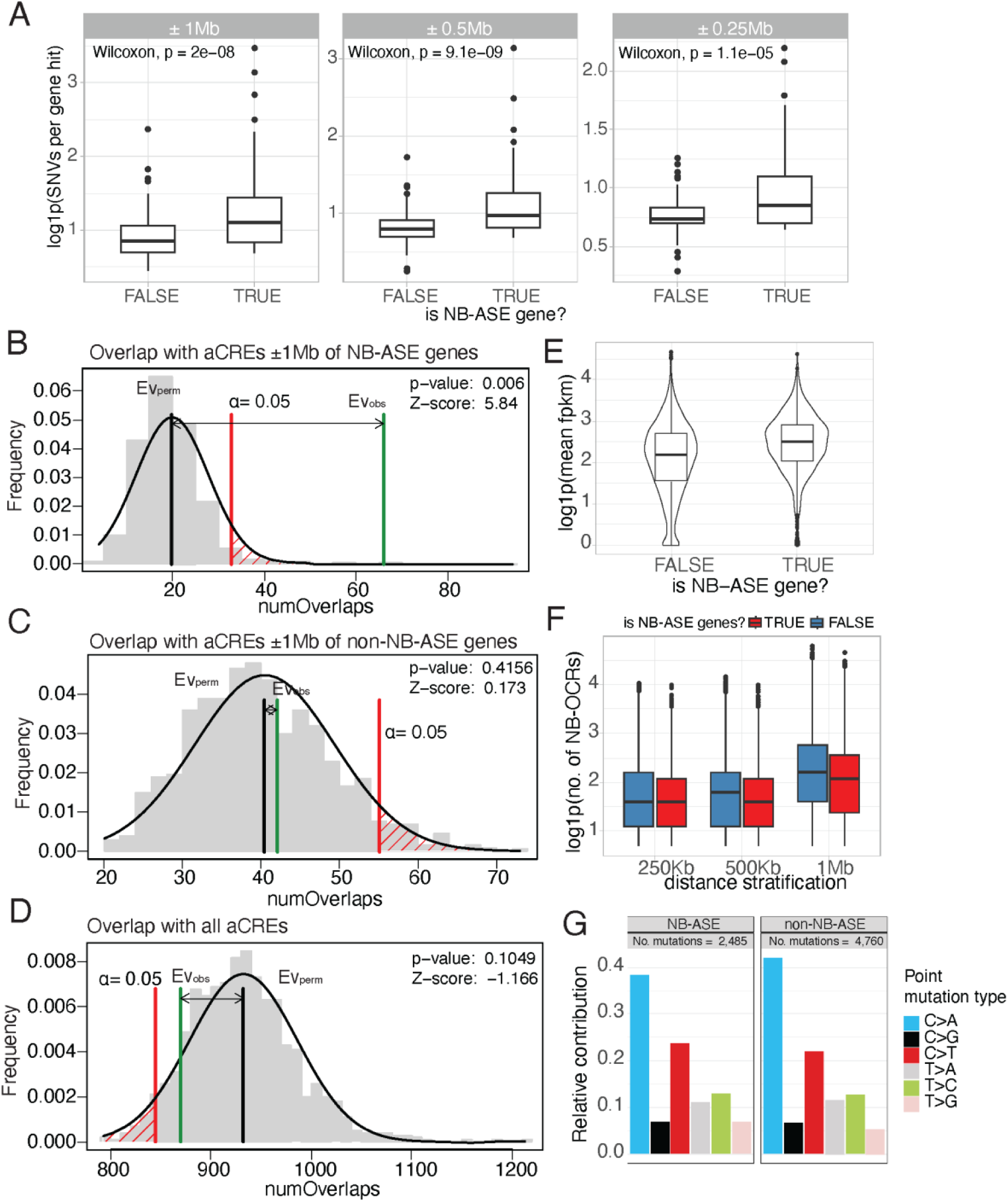
Noncoding somatic mutations are preferentially enriched in neuroblastoma-specific open chromatin regions (NB-OCRs) near NB-ASE genes, independent of regulatory density or local mutational processes. **A)** Noncoding somatic SNVdensity near NB-ASE and non-NB-ASE genes was calculated per sample and compared using a paired Wilcoxon test. **B–D)** Enrichment of noncoding somatic SNVs within neuroblastoma-specific open chromatin regions (NB-OCRs), tested against a null generated by circular randomization of SNV positions. SNVs within ±1 Mb of NB-ASE gene TSSs were significantly enriched in NB-OCRs. SNVs within ±1 Mb of genes with allelic imbalance in both neuroblastoma and normal adrenal gland showed no enrichment. All noncoding SNVs showed no enrichment. **E)** Baseline gene expression, shown as log1p(mean FPKM), for genes stratified by NB-ASE status (FALSE, non-NB-ASE; TRUE, NB-ASE). **F)** Local regulatory element density, shown as log1p(number of aCREs), within ± 250 kb, 500 kb, and 1 Mb windows around NB-ASE (red) and non-NB-ASE (blue) genes. **G)** Relative contribution of the six base-substitution classes among somatic single-nucleotide variants located near NB-ASE genes (n = 2,485 mutations) and non-NB-ASE genes (n = 4,760 mutations).

Prior studies have used epigenetic data (e.g. chromatin accessibility and histone modification) from neuroblastoma cell lines to identify functional noncoding mutations [13, 14]. However, these studies have not directly tested whether noncoding mutations are enriched within neuroblastoma specific open chromatin regions (NB-OCRs). To examine this association, we tested whether noncoding SNVs were preferentially enriched within NB-OCRs surrounding NB-ASE genes. To identify NB-OCRs, we compared chromatin accessibility profiles from neuroblastoma cell lines with human neural crest cells while controlling for MYCN amplification status, age and sex. Using a false discovery rate (FDR) ≤ 0.05 or 5% and an absolute log_2_ fold change ≥ 1.5, we identified 22,155 candidate NB-OCRs. We then assessed the overlap between noncoding SNVs and NB-OCRs relative to a null distribution generated through circular randomization of SNV positions. This approach preserves the local clustering of somatic mutations while disrupting their positional relationship with accessible chromatin.

We observed that SNVs located within ±1 Mb of transcription start sites of NB-ASE genes were significantly enriched within NB-OCRs (Figure 2B). In contrast, SNVs located within ±1 Mb of non-NB-ASE genes showed no significant enrichment (Figure 2C), nor did the full set of noncoding somatic SNVs when intersected with NB-OCRs genome-wide, without restriction to specific gene loci (Figure 2D). This enrichment was robust to window size: repeating the analysis at ±500 kb and ±250 kb yielded consistent enrichment results (Supplementary Figure S2). Together, these findings indicate that enrichment of noncoding mutations within NB-OCRs is specific to NB-ASE gene loci, rather than a general property of neuroblastoma somatic mutations or of ASE genes overall, and that this specificity holds across multiple window sizes.

Because baseline gene expression, local regulatory element density, and mutation spectrum could potentially influence cis-regulatory overlap, we examined each separately as a potential confounding factor in this enrichment. First, we observed that NB-ASE genes showed modestly higher baseline expression (log1p mean fragments per million kilobases) than non-NB-ASE genes (Figure 2E). Second, we observed that the number of NB-OCRs within ± 0.25Mb, ±0.5Mb, and ±1Mb of each gene was broadly similar between NB-ASE and non-NB-ASE genes across all window sizes (Figure 2F). Finally, to determine if the difference in enrichment patterns could be explained by differences in mutational process acting on NB-ASE and non-NB-ASE genes, we compared the relative contributions of the six base-substitution classes among somatic SNVs within ±1 MB of NB-ASE (2,485 mutations) and non-NB-ASE (4,760 mutations) genes. We observed that both groups exhibited similar mutational spectra dominated by C>A and C>T substitutions (cosine similarity = 0.98) with comparable relative contributions across all six substitution classes (Figure 2G and Supplemental Figure S3). These analyses indicate that the enrichment of somatic mutations within NB-OCRs near NB-ASE genes is not readily explained by differences in baseline gene expression, local regulatory element density, or mutational spectrum and likely reflects biologically relevant cis-regulatory disruptions.

### TFBS–Regulon Convergence Analysis: Noncoding somatic mutations near ASE genes converge on specific transcription factor programs in neuroblastoma

To functionally interpret noncoding somatic SNVs located near NB-ASE genes, we scored each variant’s predicted regulatory effect across chromatin and TF binding tracks from neuroblastoma-relevant cell lines (SK-N-SH, SK-N-DZ, SK-N-BE2C, SH-SY5Y) and human neural crest cells (hNCC) using AlphaGenome [28]. Of 2,485 noncoding SNVs evaluated, 792 (32%) had an absolute quantile-normalized AlphaGenome effect score ≥ 0.95 for at least one chromatin or TF track and were retained for downstream analysis (Supplemental Figure S4). The AlphaGenome quantile normalize score is calculated by comparing a variant’s raw predicted effect against a fixed background reference distribution of 348,126 common human SNPs from gnomAD. Of these 792 SNVs, 789 were private (occurred in only one sample), making them unsuitable for direct association testing with ASE. We therefore sought alternative ways to infer potential regulatory effects of these SNVs on ASE genes.

To this end, we came up with a novel strategy which we call as TFBS-Regulon convergence analysis. In this analysis we identified noncoding SNVs predicted to alter TFBS within ±1Mb of transcription start sites of NB-ASE genes. Then we examined whether these NB-ASE genes were preferentially enriched among strong regulatory targets of corresponding TFs. Under the null hypothesis that these SNVs do not preferentially perturb transcriptional regulation through the affected TFs, NB-ASE genes near disrupted binding sites should show no enrichment within the corresponding TF regulons. Conversely, enrichment of nearby ASE genes among targets of a specific TF would provide evidence of convergence of noncoding SNVs on that TF’s regulatory program.

To identify TFs whose predicted binding disruption at noncoding SNVs is preferentially associated with co-regulated NB-ASE genes, we first constructed a gene regulatory network from RNA-seq expression data across112 neuroblastoma tumors for a panel of 30 TFs using GENIE3 [29]. These TFs were selected because ChIP-seq data were available in at least one neuroblastoma cell line and they were included in the published AlphaGenome training set. For each TF, we then identified NB-ASE genes that harbored at least one nearby TFBS SNV (within ±1 Mb of the transcription start site) with an absolute AlphaGenome quantile score ≥ 0.95, indicating a medium to strong predicted effect on their TFBS. To test whether these genes formed unusually strong regulatory modules, we compared their mean GENIE3 edge weights against a null distribution generated by randomly sampling target genes from the same TF’s regulatory network (10,000 permutations). Of the 30 TFs tested, four show enrichment after multiple-testing correction (padj ≤ 0.1 or 10%): GATA3 (104 NB-ASE-linked target genes; padj= 0.002), ZBTB33 (106 genes; padj = 0.014), GABPA (106 genes; padj = 0.024), and JUND (104 genes; padj = 0.098) (Figure 3A). GENIE3 edge weight showed only a weak correlation with target gene baseline expression level (Spearman’s ρ = −0.023), arguing against baseline expression as a confounder of this result.

**Figure 3:**
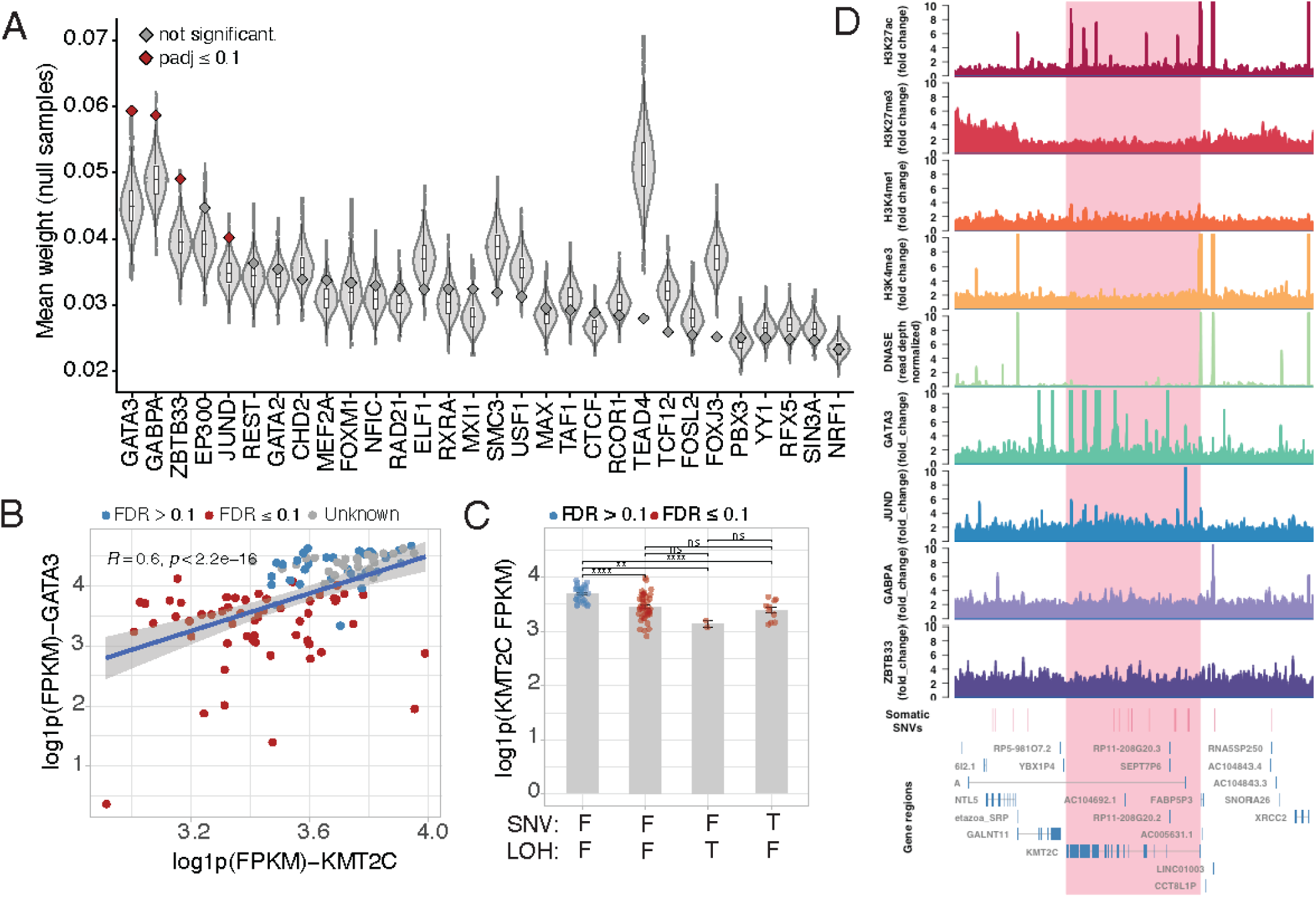
GATA3 regulon convergence and disruption of a GATA3-bound enhancer near KMT2C. **A)** Mean GENIE3 edge weight for NB-ASE genes harboring a nearby TFBS-disrupting SNV for 30 tested TFs (Red and Gray diamonds), relative to null distributions from 10,000 permutations of randomly sampled target genes from the same TF’s regulatory network. Red diamonds, TFs with significant enrichment (padj ≤ 0.1); Gray diamonds, not significant. GATA3 shows the strongest enrichment (padj = 0.002). **B)** Correlation between GATA3 and KMT2C expression (log1p FPKM) across neuroblastoma tumors, colored by KMT2C ASE FDR status (Pearson R = 0.6, p < 2.2 × 10⁻¹⁶). **C)** KMT2C expression (log1p FPKM) stratified by ASE FDR status, LOH status, and presence of a somatic noncoding SNV at the locus. Samples with LOH or SNV show significantly reduced KMT2C expression relative to non-ASE samples. LOH and SNV samples were mutually exclusive. **D)** Genome browser view of the KMT2C locus in SK-N-SH cells showing H3K27ac, H3K27me3, H3K4me1, and H3K4me3 ChIP-seq, DNase-seq accessibility, and GATA3, JUND, GABPA, and ZBTB33 ChIP-seq occupancy, with somatic SNVs and gene models below. Shaded region marks the intronic GATA3-bound enhancers that overlap somatic SNVs within this region.

Our results suggest that noncoding mutations in neuroblastoma may impact regulatory regions bound by GATA3 and play a critical role in neuroblastoma tumorigenesis. An important example of a gene that may be dysregulated by noncoding somatic mutations affecting GATA3 binding sites is KMT2C (MLL3). KMT2C encodes a large histone methyltransferase that plays a critical role in enhancer activation, cellular differentiation, and development and acts as a tumor suppressor in many adult cancers [30]. Across neuroblastoma samples, GATA3 expression was positively correlated with KMT2C expression, with samples exhibiting ASE showing reduced KMT2C expression (Figure 3B). We identified 38 somatic mutations mapping to 14 neuroblastoma samples within intronic and distal intergenic regions within ±1 Mb of the transcription start site for KMT2C. Of these, 17 SNVs mapping to 10 samples had an absolute AlphaGenome quantile score ≥ 0.95 for at least one chromatin or transcription factor binding track in neuroblastoma cell lines (Supplemental Figure S5) and 8 of these mutations mapping to 8 different samples were located within intronic regions of KMT2C. At least two of these mutations showed concordant changes in GATA3 binding and local chromatin state in SK-N-SH cells: chr7:152381189:G>A showed reduced GATA3 binding along with reduced H3K27ac and H3K4me1/2 signal, consistent with disruption of a GATA3-bound intronic enhancer element, while chr7:152377257:C>A showed the inverse pattern (Supplemental Table S5 & S6, Supplemental Figure S5). In addition to noncoding somatic mutations, we also identified two samples in which KMT2C was impacted by a loss of heterozygosity (LOH) event, suggesting that multiple genetic mechanisms cooperate to alter KMT2C expression.

To determine the effect of these mutations on bulk KMT2C expression, we next stratified neuroblastoma samples by ASE status, the presence of LOH, and the presence of noncoding mutations within intronic or distal intergenic regions of KMT2C and then examined log1p normalized KMT2C expression. Notably, ASE samples harboring either LOH or noncoding mutations in the KMT2C locus exhibited significantly lower expression compared with non-ASE samples (Figure 3C). These findings suggest that noncoding genetic variation in the KMT2C locus may contribute to reduced KMT2C expression in neuroblastoma, consistent with its known tumor suppressor function in adult tumors. Finally, we examined the chromatin features around KMT2C and observed that several of the somatic SNVs mapped to the intronic region of KMT2C overlapped GATA3 binding sites and H3K27ac in SK-N-SH NB cell lines (Figure 3D), further supporting a role for these noncoding SNVs in modulating chromatin and affecting KMT2C expression.

## DISCUSSION

Comprehensive sequencing studies of high-risk neuroblastoma have identified relatively few clinically actionable protein-coding mutations at diagnosis, motivating interest in noncoding regulatory variation as a contributor to disease pathogenesis [2]. Here, we present an integrative framework that combines ASE, chromatin accessibility profiling, and variant-effect prediction to identify candidate noncoding regulatory mutations and link them to their putative target genes and downstream transcriptional consequences in neuroblastoma.

Applying this framework to 112 neuroblastoma patients, we identified 1,363 NB-ASE genes. These genes were enriched for chromatin regulatory and neurodevelopmental processes and showed a strong association with the adrenergic transcriptional program, supporting their relevance to neuroblastoma biology. Investigation of the underlying genetic basis of NB-ASE patterns revealed that although most events were associated with SCNA, a subset occurred within copy-number-neutral regions. These SCNA-independent NB-ASE genes were significantly enriched for noncoding somatic mutations within NB-OCRs mapping to their intronic and distal intergenic regions, implicating regulatory mutations as a distinct mechanism of gene dysregulation.

These findings extend prior work in two important ways. Prior studies have demonstrated that somatic mutations within neuroblastoma-specific open chromatin impact DNA binding sites for TFs implicated in cell proliferation and chromatin regulation and converge on genes involved in embryonic development and immune response [13, 14]. However, both studies rely on mutation recurrence — either across binding sites for a given TF or across patients at a given regulatory element — as their primary evidence, which is intrinsically underpowered to detect noncoding drivers that are rare or private to individual tumors. Second, neither prior study accounted for SCNA, which are pervasive in neuroblastoma and can directly alter local gene dosage independent of any regulatory mutation. Because neither study excluded or adjusted for SCNA status when comparing gene expression between mutated and non-mutated samples, it remains possible that some previously reported associations between noncoding mutation and target-gene expression reflect co-occurring copy-number alterations rather than a direct regulatory effect.

Our framework directly addresses both limitations. First, by using ASE as a functional, patient-level readout of cis-regulatory disruption, we can nominate candidate regulatory drivers even when the underlying mutations are largely private. Second, by explicitly partitioning ASE into SCNA-associated and SCNA-independent categories, we isolate a subset of allelic imbalance that arises independently of chromosomal gain or loss and is specifically linked to nearby noncoding mutations in accessible chromatin.

Our TFBS-regulon convergence analysis also extends previous approaches for interpreting noncoding somatic mutations in neuroblastoma. Prior studies in neuroblastoma have primarily focused on testing for enrichment of somatic mutations within TFBS within mutated regulatory regions and assigning target genes based on nearest gene assignment. In contrast, we take a network approach where we construct an expression-derived TF regulatory network and test whether NB-ASE genes harboring nearby TFBS mutations are enriched within the corresponding TF’s regulon. To our knowledge, this represents a novel framework for interpreting the functional consequences of noncoding mutations in neuroblastoma. Using this approach, we found that NB-ASE genes carrying mutations predicted to disrupt GATA3 binding sites in their regulatory regions are significantly enriched among members of the GATA3 regulon. These findings suggest that noncoding mutations in neuroblastoma converge on the GATA3 transcriptional network. GATA3 is a component of the transcriptional regulatory circuitry that maintains neuroblastoma cell identity, acting alongside crucial TFs like MYCN, PHOX2B, HAND2, ISL1, and TBX2 [31]. This observation suggest that the noncoding mutations may contribute to NB pathogenesis by perturbing the core regulatory circuitry in neuroblastoma.

Finally, our study also has several limitations. Most importantly, the variant-to-function relationships reported here rely on correlative genomic evidence rather than direct experimental validation. Future studies should prioritize experimental validation of candidate variants using approaches such as CRISPR-based perturbation and reporter assays, as well as integration of larger cohorts and chromatin conformation data to strengthen variant-to-gene assignments. Despite these limitations, our study demonstrates the complexity of neuroblastoma genomes and provides a comprehensive list of noncoding SNV-gene pairs for functional analysis in future studies.

## METHODS

### Patient Dataset

For this study, we integrated whole genome sequencing (WGS) and RNA sequencing datasets from Pediatric Cancer Genome Program (PCGP; N = 73) and Kids First Pediatric Research Program (Kids First; N= 83). Because paired normal tissues were not included in transcriptional profiling of the PCGP and Kids First dataset, we also constructed a reference panel of normals for ASE analysis using 220 phenotypically normal adrenal gland samples from the Genotype-Tissue Expression project (GTEx; RRID: SCR_013042).

### Allele-specific copy number analysis

Allele-specific copy number, tumor purity, and tumor ploidy was estimated from WGS data using FACETS (RRID:SCR_026264) [16]. Samples with a tumor purity estimate below 30%, at which copy-number calls made using FACETS become unreliable, were excluded from analysis. Data analysis and visualization was performed in R version 4.5.2 (RRID: SCR_001905). Segments were classified from FACETS EM estimates relative to each tumor’s ploidy: gain (tcn.em > round(ploidy)), loss (tcn.em < round(ploidy)), copy-neutral LOH (tcn.em = round(ploidy), lcn.em = 0), and copy-neutral (tcn.em = round(ploidy), lcn.em = round(ploidy)/2 as specified by FACETS documentation.

### Comparing ASE profiles of neuroblastoma and adrenal gland samples

To generate allele-specific expression (ASE) profiles, we first identified heterozygous variants located within gene exons. For neuroblastoma samples, WGS reads were aligned to the human reference genome (GRCh38) using BWA-MEM (RRID: SCR_010910) and nonduplicated reads with a mapping quality score (MAPQ) ≥ 30 were retained using SAMtools (v1.9) (RRID: SCR_002105)[32]. Samples were then genotyped and high-quality heterozygous variants were identified using the Genome Analysis Toolkit (GATK4; RRID: SCR_001876) [33]. For GTEx samples, heterozygous variants were obtained from the GTEx WGS variant call set. Heterozygous exonic variants from both datasets were annotated using ANNOVAR (RRID:SCR_012821) [34] and subsequently integrated with RNA-seq data for ASE analysis.

RNA-seq reads from neuroblastoma and GTEx samples were aligned to the human reference genome (GRCh38) using STAR (v2.7.3a) (RRID: SCR_004463) [35], and reads with MAPQ ≥ 20 were retained using SAMtools (v1.9) [32]. Reads overlapping heterozygous variants were identified, and reference mapping bias was corrected using WASP to obtain accurate allele-specific read counts [36]. Heterozygous variants covered by at least 10 reads were retained for ASE analysis. Gene-level ASE was estimated using our previously published algorithm, which integrates information across multiple heterozygous sites mapping to genes while accounting for genotyping error, sequencing error, overdispersion of read counts, and variant phasing [11]. Neuroblastoma-specific ASE (NB-ASE) genes were defined as genes that were testable for ASE in at least 10 neuroblastoma and 10 adrenal gland samples and exhibited significant ASE (FDR ≤ 0.1) in at least 3 neuroblastoma samples and in no more than 1 adrenal gland sample, similar to out published study [37]. Data was visualized in R version 4.5.2 (RRID: SCR_001905) using the ggplot2 package (version 4.0.1; RRID: SCR_014601).

### Gene Ontology Analysis

Gene Ontology (GO) Biological Process term enrichment was performed in R v4.5.2. NB-ASE genes were tested against a background gene universe of all ASE genes identified in the neuroblastoma cohort. A binary gene list (target vs. universe) was constructed and analyzed using the topGO package (RRID:SCR_014798). GO annotations and mapping of gene symbols to Entrez identifiers retrieved using the org.Hs.eg.db annotation package in R (RRID:SCR_024739). GO terms were tested using the “weight01” algorithm which account for GO graph topology and dependency structure between terms. A node size threshold of 10 genes was used to exclude very small, poorly powered terms. Statistical significance was assessed using Fisher’s exact test, and the top 25 enriched terms were retained, ranked by weighted Fisher p-value. Enrichment results were visualized as a dot plot using ggplot2 with dot size representing the number of significant genes in each term and color representing −log10 (p-value).

### Processing neuroblastoma cell line chromatin accessibility data

Assay for Transposase-Accessible Chromatin with high-throughput sequencing (ATAC-seq) data for neuroblastoma cell lines and human neural crest cells were obtained from GSE138315 and GSE108517 [38, 39]. Sequencing reads were aligned to the GRCh38 reference genome using BWA-MEM (v0.7.17) [40] and non-duplicated reads with a MAPQ ≥ 20 were retained using SAMtools (v1.9). Narrow peaks were identified with MACS2 (v2.1.1.2) (RRID: SCR_013291) using parameters recommended by the ENCODE project (RRID:SCR_015482) (P ≤ 0.01). To compare chromatin accessibility profiles across neuroblastoma cell lines and between neuroblastoma cell lines and human neural crest cells, the GRCh38 reference genome was partitioned into non-overlapping 1Kb bins. Bins overlapping peaks identified in at least one neuroblastoma cell line were retained and read counts for these bins were obtained using ChrAccR (v0.9.21). Differential chromatin accessibility to identify neuroblastoma-specific open chromatin regions between neuroblastoma cell lines and human neural crest cells was performed using DESeq2 (v1.41.13) while controlling for MYCN status, age, and sex of neuroblastoma cell lines.

### Identifying somatic mutation neuroblastoma samples

We assessed the enrichment of noncoding SNVs within regulatory regions of NB-ASE genes and prioritized candidates by evaluating their impact on transcriptional factor binding sites (TFBS) and chromatin features. We identified somatic SNVs by comparing WGS data from tumor and matched normal samples using Mutect2 (RRID:SCR_026692) [41]. For each tumor, somatic mutation was filtered for high-quality calls using the recommended GATK Best Practices pipeline: cross-sample contamination was estimated using GetPileupSummaries and CalculateContamination (GATK4; RRID:SCR_001876) against common SNP sites from ExAC, and candidate variants were filtered using FilterMutectCalls with the resulting contamination estimates. These variants were further filtered using DP >10 and then mapped to genomic regions using ANNOVAR [34].

### Somatic mutation filtering and enrichment test

Our previously published study demonstrated that somatic copy number alterations (SCNA) and loss of heterozygosity (LOH) are a major driver of ASE in neuroblastoma [11], which could confound the attribution of ASE signal to noncoding SNVs. To address this limitation, we restricted our downstream analysis to ASE genes and SNVs overlapping copy neutral regions in each tumor sample. SCNA and LOH regions were identified using FACETS as described previously [16]. Additionally, we also removed SNVs that intersected with unmappable regions of the human genome for reference genome GRCh38. Somatic SNVs were then restricted to noncoding classes (intergenic, intronic, upstream and downstream annotations). Three variant sets were evaluated: (i) SNVs located within ±1 Mb of an NB-ASE gene (“case”); (ii) SNVs located within ±1 Mb of a testable gene not classified as NB-ASE (“control”); and (iii) all noncoding somatic SNVs in the cohort (“all”).

Enrichment analysis for noncoding SNVs within NB-OCRs was performed using the regioneR(v1.42.0) (RRID:SCR_028251) [42]. For each test, SNV positions were randomized using circular randomization (circularRandomizeRegions), which shifts variant positions along each chromosome while preserving their relative spacing and thereby the local clustering structure of somatic mutations; NB-OCRs were held fixed. Overlap was quantified as the number of SNVs overlapping at least one accessible region (numOverlaps with count.once = TRUE). Randomized regions were required not to overlap the mask (max.mask.overlap = 0). Each test comprised 1,000 permutations, and significance was assessed by a two-sided Z-test against the permuted distribution. Observed overlap counts, permuted means and standard deviations, Z-scores, and permutation P values are reported for each variant set.

### Mutational spectrum analysis of SNVs near case and control genes

SNVs located within ±1 Mb of ASE genes were partitioned into two sets: those near NB-ASE (case) genes, and those near a matched set of non— NB-ASE control genes. Variant coordinates, reference alleles, and alternate alleles were parsed and converted to GRanges objects using the GenomicRanges package (RRID:SCR_017051) in R, with chromosome naming and lengths standardized to the UCSC hg38 human reference genome build (BSgenome.Hsapiens.UCSC.hg38). Trinucleotide-context mutation count matrices (96-channel SNV classification) were generated for the case and control variant sets using the MutationalPatterns package [43]. Mutational spectra were visualized as 96-channel trinucleotide profiles and summarized further into six base-substitution classes for direct comparison between case and control sets. Similarity between the case and control mutational profiles was quantified using cosine similarity.

### Analyzing variant effect on chromatin accessibility and histone modification

We used AlphaGenome to predict allele-specific effects of somatic SNVs on transcription factor binding sites, chromatin accessibility, and histone modification profile of neuroblastoma cell lines and human neural crest cells [28]. For each variant, matched sequence windows containing either the reference or alternate allele were evaluated across all assay tracks using batch variant scoring pipeline described in the AlphaGenome tutorial (https://www.alphagenomedocs.com/colabs/batch_variant_scoring.html). AlphaGenome outputs quantile scores which is calculated by comparing a variant’s raw predicted effect against a fixed background reference distribution of 348,126 common human SNPs from gnomAD. We used a threshold of greater than or equal to the absolute value of 0.95 to identify putative functional variants. Figures were generated in R version 4.5.2 using ggplot2 (v4.0.1) and ComplexHeatmap (v2.26.0) (RRID: SCR_017270).

### Co-expression network inference and TF–target enrichment

Read counts from PCGP and Kids First RNA-seq data were obtained using the featureCounts (RRID: SCR_012919) function from the Subread (RRID:SCR_009803) (v2.0.6) and were normalized using fragments per kilobase per million (FPKM) using DESeq2 (RRID:SCR_015687) (v1.41.13). FPKM normalized expression from the PCGP and Kids First patients were merged and genes with zero variance and mean FPKM ≤1 were excluded. The filtered FPKM matrix was log1p transformed before co-expression network analysis.

A regulatory network for transcription factors was inferred using a random-forest-based inference method called GENIE3 (RRID:SCR_000217) (v1.32.0) [29]. Briefly, GENIE3 decomposes the prediction of a regulatory network between p genes into p different regression problems. In each of the regression problems, the expression pattern of one of the genes (transcription factor) is predicted from the expression patterns of all the other genes (input genes), using tree-based ensemble methods Random Forests or Extra-Trees. Pairwise transcription factor and gene target importance scores (edge weights) were extracted for downstream analyses.

To determine whether NB-ASE genes harboring at least one predicted transcription factor binding site– altering SNV within ±1 Mb of their transcription start site were more likely than expected by chance to belong to the corresponding transcription factor regulon, we compared their mean GENIE3 edge weight with a null distribution generated from 10,000 random permutations. In each permutation, a gene set of equal size was sampled without replacement from all expressed genes, and the mean GENIE3 edge weight between the transcription factor and the sampled genes was calculated. Statistical significance was assessed using an empirical permutation P value, defined as the proportion of permuted mean edge weights greater than or equal to the observed mean. Using this approach, we tested enrichment of NB-ASE genes among predicted regulon members for transcription factors that were expressed in our neuroblastoma cohort and had at least 100 SNVs predicted to impact its DNA binding site. All figures were generated in R version 4.5.2 using ggplot2 (v4.0.1).

### Ethics statement

This study is a secondary analysis of previously collected de-identified human genomic and transcriptomic data. WGS and RNA-seq data from the Pediatric Cancer Genome Project and the Kids First Pediatric Research Program were obtained under approved data access requests (dbGaP: phs001436.v1.p1.c1), and GTEx data were obtained under dbGaP accession phs000424. No new human participants were recruited, and no new primary human-subject data were generated for this study. Under the U.S. Common Rule (45 CFR 46), secondary analysis of de-identified data does not constitute human subjects’ research.

## Declaration of interests

The authors declare no potential conflicts of interest.

## Acknowledgements

We thank the Pediatric Cancer Genome Project for access to data. This work was supported by the UAB Impact Fund and UAB Genetics Research Division. During the preparation of this work the author(s) used Anthropic Claude in order to edit grammar, spelling, and sentence construction. After using these tool/service, the author(s) reviewed and edited the content as needed and take(s) full responsibility for the content of the publication

## Author Contributions

B.J performed all analysis and wrote the initial draft, G.S and A.R helped B.J with analysis, A.S. conceptualized and supervised the project and wrote the paper.

## Data and code availability

WGS and RNA-seq data from the Pediatric Cancer Genome Project are available through St. Jude Cloud under controlled access [44]. Kids First Pediatric Research Program data are available through the Kids First Data Resource Portal (dbGaP: phs001436.v1.p1.c1; RRID: SCR_002709) under controlled access. Genotype and RNA sequencing data from GTEx are available through dbGaP (phs000424) under controlled access. Neuroblastoma cell line and human neural crest ATAC-seq data is available through gene expression omnibus (RRID: SCR_005012) (GSE138315 and GSE108517). SK-N-SH NB cell line epigenetic data are available through the ENCODE portal [45] with the following identifiers: ENCFF120SAA, ENCFF105KVL, ENCFF344RYH, ENCFF012OQH, ENCFF840FTR, ENCFF755FHH, ENCFF262UEH, ENCFF224TVD, ENCFF280RMA. Copy number output for the PCGP and Kids First cohort is provided in supplementary table S3; Code for allele-specific expression analysis is available through CodeOcean (https://codeocean.com/capsule/8770085/tree) and was part of a published paper[11]. This study did not generate new sequencing datasets.

## Notes

### Competing Interest Statement

The authors have declared no competing interest.

### Summary of Updates

For this manuscript, all analysis have been updated to include more robust controls datasets. This have resulted in significant changes in final results.

