## supplemental figures for "Noncoding regulatory mutations contribute to the aberrant gene expression profile of neuroblastomas"

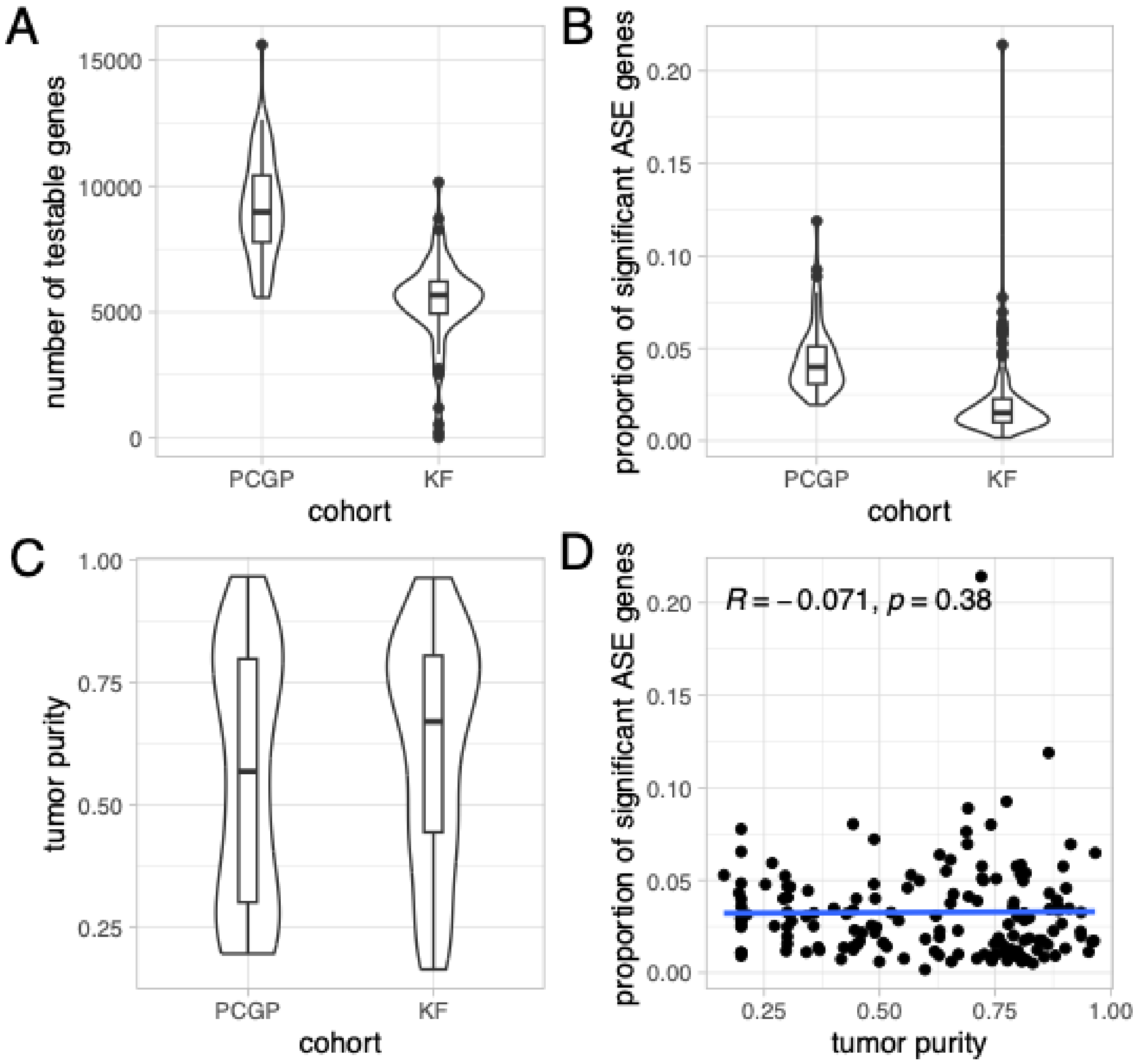


**Supplemental Figure 1. Cohort-level quality-control metrics for ASE testing in the PCGP and Kids First (KF) neuroblastoma cohorts. A)** Number of genes testable for allele-specific expression (ASE) per tumor sample, stratified by cohort. PCGP samples had a higher median number of testable genes than KF samples. **B)** Proportion of testable genes exhibiting significant ASE (FDR ≤ 0.1 or 10%) per tumor sample stratified by cohort. **C)** FACETS-derived tumor purity estimates per sample, stratified by cohort. **D)** Relationship between tumor purity and the proportion of genes exhibiting significant ASE per sample across both cohorts combined. No significant association was observed (Pearson's R = −0.071, P = 0.38), indicating that ASE detection was largely robust to variation in tumor purity within this dataset.

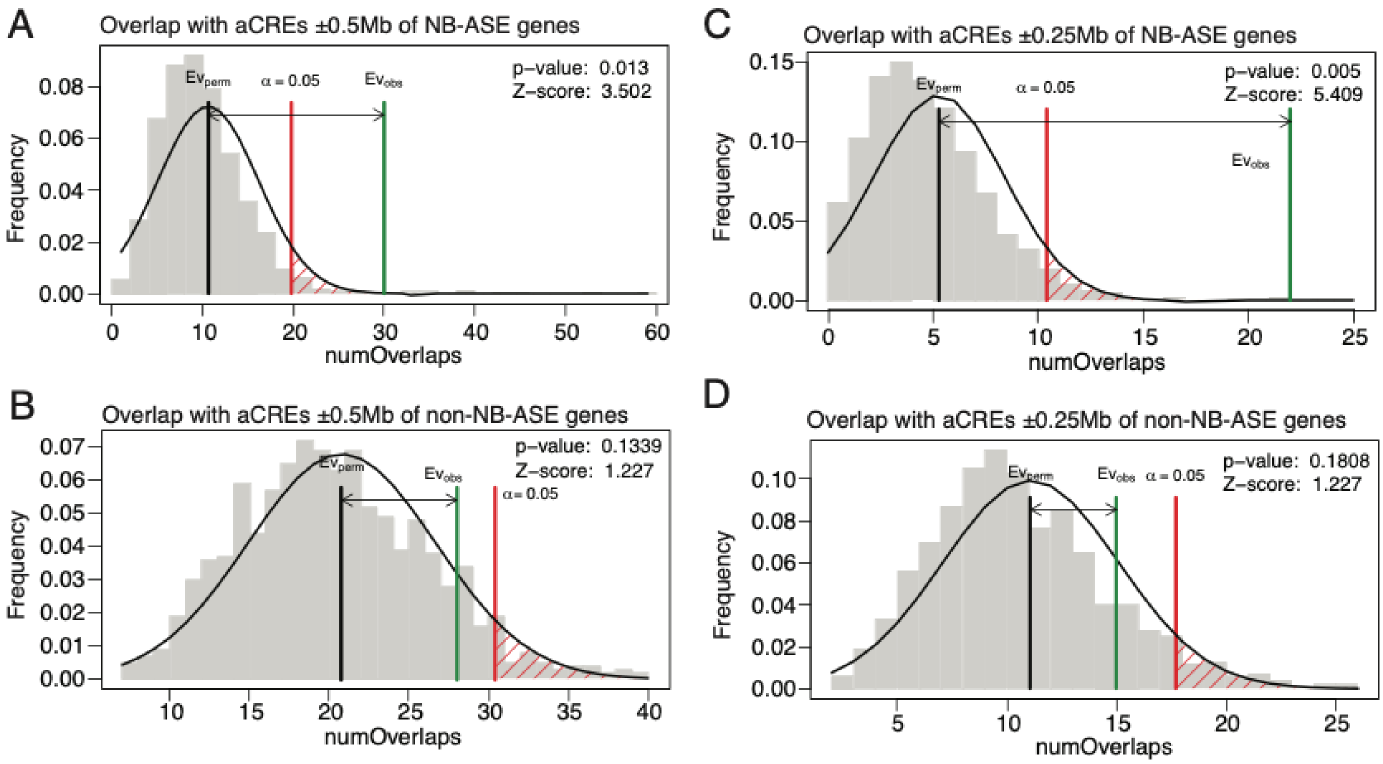


**Supplemental Figure 2. Enrichment of noncoding mutations within neuroblastoma-specific open chromatin regions (NB-OCRs) near NB-ASE genes is robust across smaller genomic window sizes.** Permutation-based enrichment analysis of noncoding somatic SNV overlap with NB-OCRs, restricted to windows of ±500 kb (A & B) and ±250 kb (C & D) around the transcription start sites of NB-ASE and non-NB-ASE genes (Figure 2). Null distributions (grey histograms) were generated by circular randomization of SNV positions, which preserves local mutation clustering while disrupting positional relationships with NB-OCRs. In each panel, the black vertical line indicates the mean of the null distribution, the red line indicates the significance threshold ($\alpha$ = 0.05), and the green line indicates the observed number of overlaps. These results confirm that enrichment of noncoding mutation within NB-OCRs is specific to NB-ASE gene loci and is not an artifact of the ±1 Mb window used in the primary analysis.

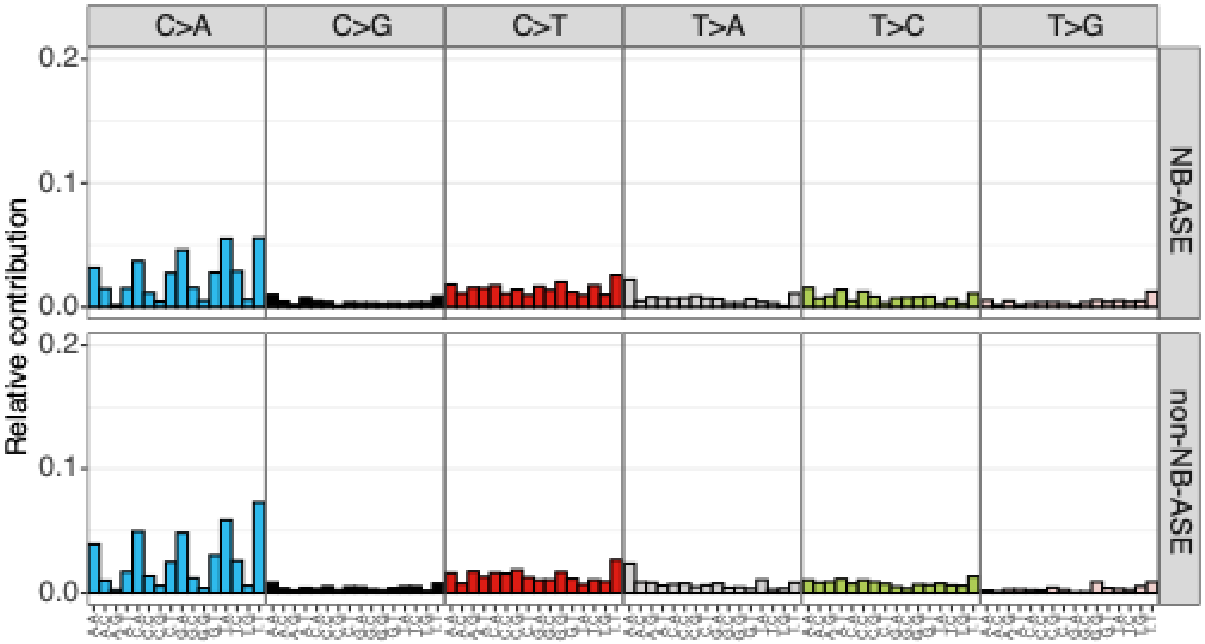


**Supplemental Figure S3. Trinucleotide mutational context profiles are similar near NB-ASE and non-NB-ASE genes.** 96-trinucleotide mutational profiles for somatic single-nucleotide variants located near NB-ASE genes (2,637 mutations) and non-NB-ASE genes (4,646 mutations), generated using MutationalPatterns. For each of the six base-substitution types (C>A, C>G, C>T, T>A, T>C, T>G), the relative contribution of each of the 16 possible trinucleotide contexts (5′ and 3′ flanking bases) is shown. The two mutational spectra were highly similar (cosine similarity = 0.98) indicating that local mutational processes do not differ substantially between SNVs near NB-ASE and non-NB-ASE genes and are therefore unlikely to explain the enrichment of noncoding mutations within NB-OCRs specifically at NB-ASE gene loci (Figure 2G).

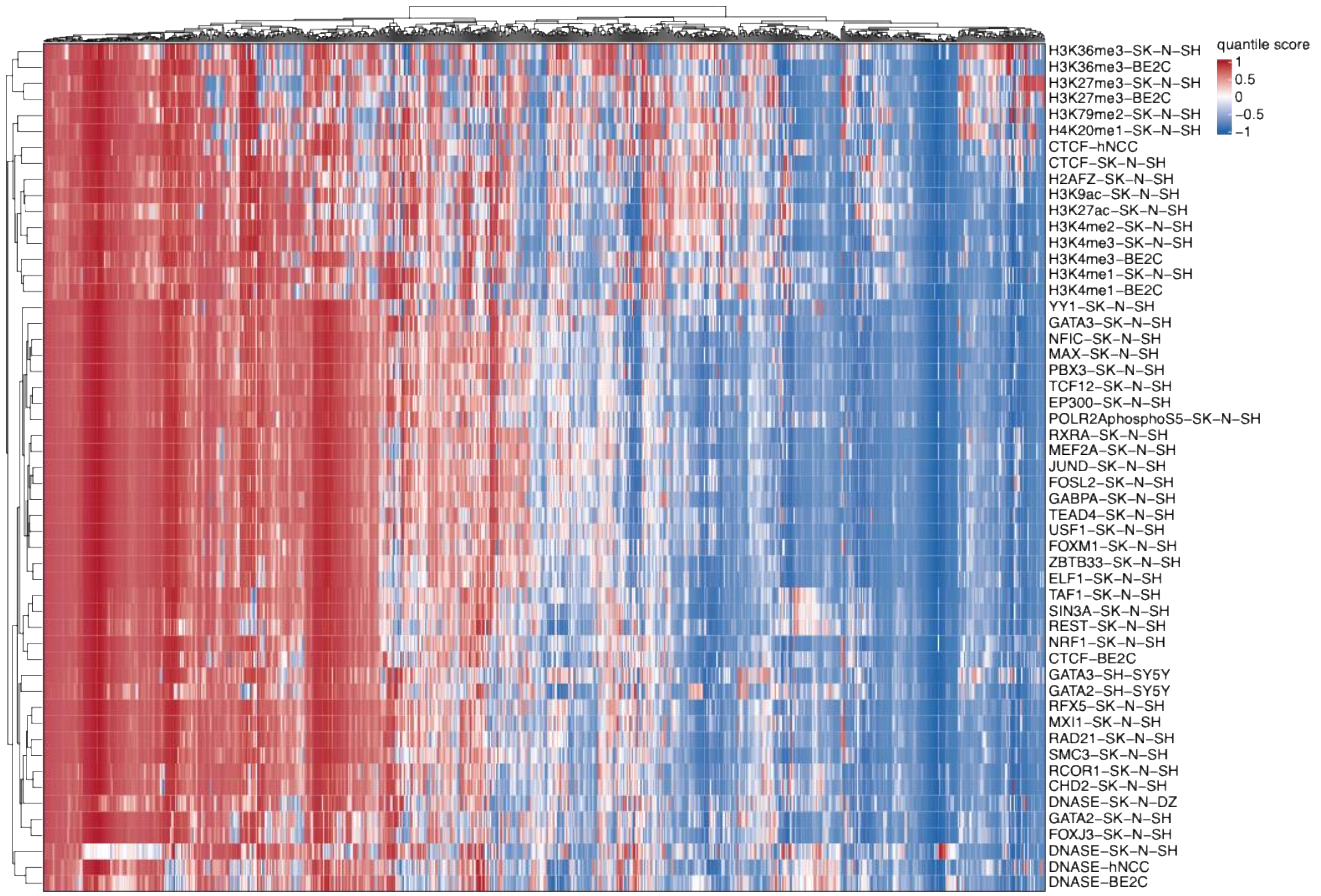


**Supplemental Figure S4.** AlphaGenome-based functional scoring classifies noncoding SNVs near NB-ASE genes. Heatmap of AlphaGenomequantile scores for 792 noncoding SNVs near NB-ASE genes (columns) with a significant predicted regulatory effect (|quantile score| ≥ 0.95) in at least one of chromatin and transcription factor (TF) binding tracks (rows) from neuroblastoma cell lines (SK-N-SH, SK-N-DZ, BE2C, SH-SY5Y) and human neural crest cells (hNCC). Color indicates quantile score, from −1 (blue) to 1 (red).

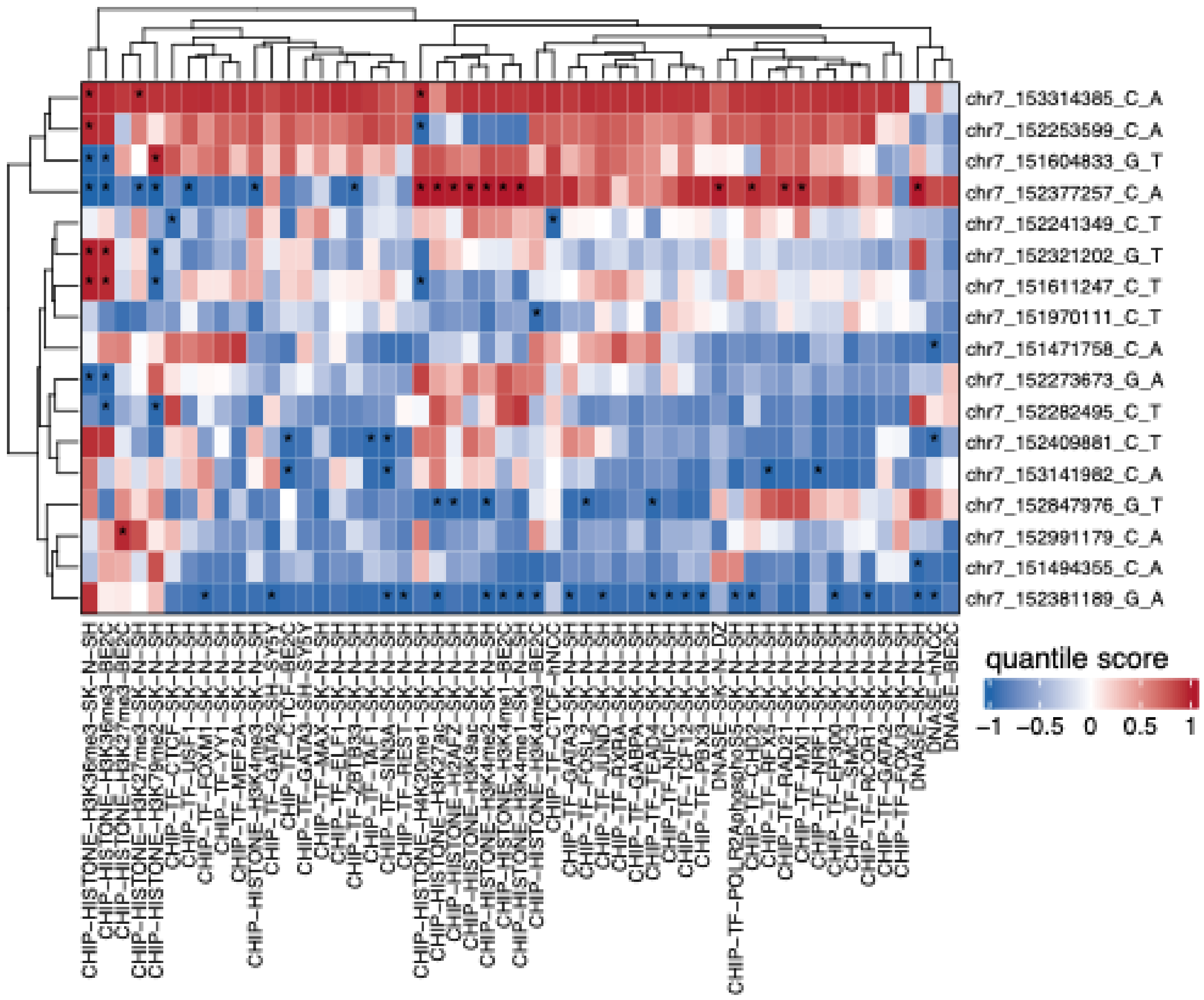


**Supplemental Figure S5:** AlphaGenome based functional scoring classifies noncoding SNVs near KMT2C. Heatmap of AlphaGenome quantile scores for 17 noncoding SNVs near NB-ASE genes (columns) with a significant predicted regulatory effect (|quantile score| ≥ 0.95; indicated by *) in at least one of 47 chromatin and transcription factor (TF) binding tracks (rows) from neuroblastoma cell lines (SK-N-SH, SK-N-DZ, SK-N-BE2C, SH-SY5Y) and human neural crest cells (hNCC).
