## supplemental table legends for "Noncoding regulatory mutations contribute to the aberrant gene expression profile of neuroblastomas"

**Supplemental Tables.**

**Supplemental Table S1.** Allele specific expression (ASE) profile of neuroblastoma tumor samples from Pediatric Cancer Genome Project (PCGP) and Kids First Pediatric Research Program (KF).

**Supplemental Table S2.** For each tumor sample, we used FACETS to estimate allele-specific copy number from matched whole genome sequencing data. This table reports the sample-level statistics provided by FACETS. Columns report: sample, sample identifier; purity, estimated tumor cellular fraction; ploidy, estimated average tumor ploidy; dipLogR, the log-ratio value corresponding to the diploid copy-number state, used as a reference point for calling absolute copy number; flags, FACETS quality flags (e.g., low-purity or unreliable-estimate warnings; blank indicates no flag raised); n_seg, total number of copy-number segments called; n_loh, number of segments exhibiting loss of heterozygosity (LOH); and loh_frac, the fraction of the genome affected by LOH.

**Supplemental Table S3.** This table contains the segmental allele-specific copy number calls for tumor samples outputted by FACETS.

**Supplemental Table S4.** The table summarizes the total number of samples testable for allele-specific expression (ASE) and the number of samples exhibiting significant ASE (FDR $\leq 0.1$) for each gene. Genes were classified as neuroblastoma-specific ASE genes (hard_filter = TRUE) if they met all of the following criteria: (i) were testable for ASE in at least 10 neuroblastoma and adrenal gland samples, (ii) exhibited significant ASE in at least 3 neuroblastoma samples, and (iii) exhibited significant ASE in no more than 1 adrenal gland sample.

**Supplemental Table S5.** This table provides a comprehensive catalog of all single-nucleotide variants (SNVs) located within ±1 Mb of transcription start sites of neuroblastoma-specific ASE genes and mapped to copy-neutral genomic regions in each sample. The table includes genomic coordinates, allelic information, and sample-level annotations for variants associated with neuroblastoma-specific ASE genes.

**Supplemental Table S6.** This table provides the raw prediction scores and quantile scores from AlphaGenome based variant effect predictions for SNVs located with ±1 Mb of transcription start sites of neuroblastoma specific ASE genes within copy neutral regions.

**Supplemental Table S7.** For each transcription factor (TF), n_nbase is the number of NB-ASE genes with a nearby predicted TF-binding-disrupting SNV (AlphaGenome |quantile score| ≥ 0.95) used as that TF's target set; obs_mean is the observed mean GENIE3 edge weight among those genes; pval is the empirical permutation P-value (10,000 permutations, null generated by resampling from the TF's full target set); padj is the FDR-corrected P-value across all 30 TFs. Four TFs — GATA3, ZBTB33, GABPA, JUND — were significant at padj ≤ 0.1 (Figure 4A).
